# Widespread nociceptive maps in the human neonatal somatosensory cortex

**DOI:** 10.1101/2021.07.29.454164

**Authors:** Laura Jones, Madeleine Verriotis, Robert J. Cooper, Maria Pureza Laudiano-Dray, Mohammed Rupawala, Judith Meek, Lorenzo Fabrizi, Maria Fitzgerald

**Affiliations:** Department of Neuroscience, Physiology & Pharmacology, University College London, London, WC1E 6BT, UK; DOT-HUB, Department of Medical Physics & Biomedical Engineering, University College London, London, WC1E 6BT, UK; Elizabeth Garrett Anderson Obstetric Wing, University College London Hospitals, London, WC1E 6DB, UK

## Abstract

Topographic cortical maps are essential for spatial localisation of sensory stimulation and generation of appropriate task-related motor responses. Somatosensation and nociception are finely mapped and aligned in the adult somatosensory (S1) cortex, but in infancy, when pain behaviour is disorganised and poorly directed, nociceptive maps may be less refined. We compared the topographic pattern of S1 activation following noxious (clinically required heel lance) and innocuous (touch) mechanical stimulation of the same skin region in newborn infants (n=32) using multi-optode functional near-infrared spectroscopy (fNIRS). Signal to noise ratio and overall activation area did not differ with stimulus modality. Within S1 cortex, touch and lance of the heel elicit localised, partially overlapping increases in oxygenated haemoglobin (HbO), but while touch activation was restricted to the heel area, lance activation extended into cortical hand regions. The data reveals a widespread cortical nociceptive map in infant S1, consistent with their poorly directed pain behaviour.

## Introduction

Somatotopically organised cortical maps of activity evoked by innocuous or noxious mechanical stimulation allow us to localise our sense of touch or pain (Penfield and Boldrey, 1937; Harding-Forrester and Feldman, 2018), and may also convey computational advantages in the relay of afferent information to higher brain areas (Thivierge and Marcus, 2007). In adults, overlapping regions are involved in the cortical processing of noxious and innocuous mechanical stimulation (Kenshalo et al., 2000; Lui et al., 2008) and detailed fMRI analysis reveals a fine-grained somatotopy for nociceptive inputs in primary somatosensory cortex (SI) that are aligned with activation maps following tactile stimuli, suggesting comparable cortical representations for mechanoreceptive and nociceptive signals (Mancini et al., 2012).

A whole-body topographical map of innocuous mechanical stimulation develops in the sensorimotor cortices over the early postnatal period in rats, which represent the human final gestational trimester (Seelke et al., 2012). Distinct representations of the hands and feet can be observed from 31 weeks using fMRI (Dall’Orso et al., 2018), becoming increasingly localised by term age (Allievi et al., 2016). While haemodynamic responses to a clinically-required heel lance have been recorded from 28 weeks using functional near-infrared spectroscopy (Slater et al., 2006) and can be distinguished from innocuous mechanical evoked brain activity in EEG recordings from 34-35 weeks (Fabrizi et al., 2011), the source of this activity and topographic representation of these two modalities have not been mapped, or their alignment established, in the infant cortex.

Infant pain behaviour is exaggerated and disorganised in newborn rodents and human infants (Fitzgerald, 2005, 2015; Cornelissen et al., 2013). Poor spatial tuning of nociceptive reflexes and receptive fields is a feature of the developing somatosensory system, followed by the emergence of adult organisation through activity-dependent refinement of synaptic connections (Beggs et al., 2002; Schouenborg, 2008; Koch and Fitzgerald, 2013). We hypothesised that this developmental process is reflected in ascending nociceptive signals to SI, leading to widespread cortical activation and poor spatial localisation of noxious events in early life.

To test this hypothesis, we used multioptode functional near-infrared spectroscopy (fNIRS) to map nociceptive and innocuous mechanoreceptive activity across the infant sensorimotor cortex. fNIRS is a non-invasive measure of cerebral haemodynamic changes which can be performed at the bedside, using skin-to skin holding in a naturalistic hospital setting during clinically required procedures. Using the temporal and spatial profiles of haemodynamic responses to a noxious skin lance and an innocuous touch of the hand and the heel, we show that haemodynamic activity elicited by noxious and innocuous mechanical stimulus have partially overlapping topographies in the human infant S1 cortex but that the two maps are not aligned. Noxious stimulation of the heel in the newborn evokes a more widespread cortical activation than innocuous stimulation, that extends into inferior regions of S1, normally associated with representation of the hand.

## Results

### Hand and heel touch evoked activity is somatotopically organised in the newborn infant S1 cortex

We first established the cortical topography of touch activation in newborn infants by mapping the extent of activation in the contralateral somatosensory (S1) cortex following innocuous mechanical stimulation (touch) of the hand and of the heel. **Figure 1a** and **1b** show a significant and localised increase in average [HbO] in contralateral optode channels following touch of each body area (n=11, hand touch; n=16 heel touch). Touch stimulation of the hand elicited significant increases in five channels, with a maximum change (0.31 μM at 16.9 s post-stimulus) at the channel corresponding to the FCC3 position of the 10:5 placement system (Figure 4), while touch of the foot elicited significant increases in six channels, with a maximum change (0.30 μM at 15.8 s) at the channel corresponding to the CPP1 10:5 position. The somatotopically localised increases in [HbO] were accompanied by a widespread decrease in [Hb] over the whole peri-rolandic area (hand: significant decreases in eight channels [peak change: -0.21 μM at 17.7 s]; foot: significant decreases in nine channels, including control channel [peak change: -0.20 μM at 11.2 s]). An inverse response (significant decrease in [HbO], significant increase in [Hb], or both) was mostly restricted to channels surrounding the hand and foot areas of the S1, respectively. Individual channel data is shown in **Figure 1 – Source Data 1**.

**Figure 1.**
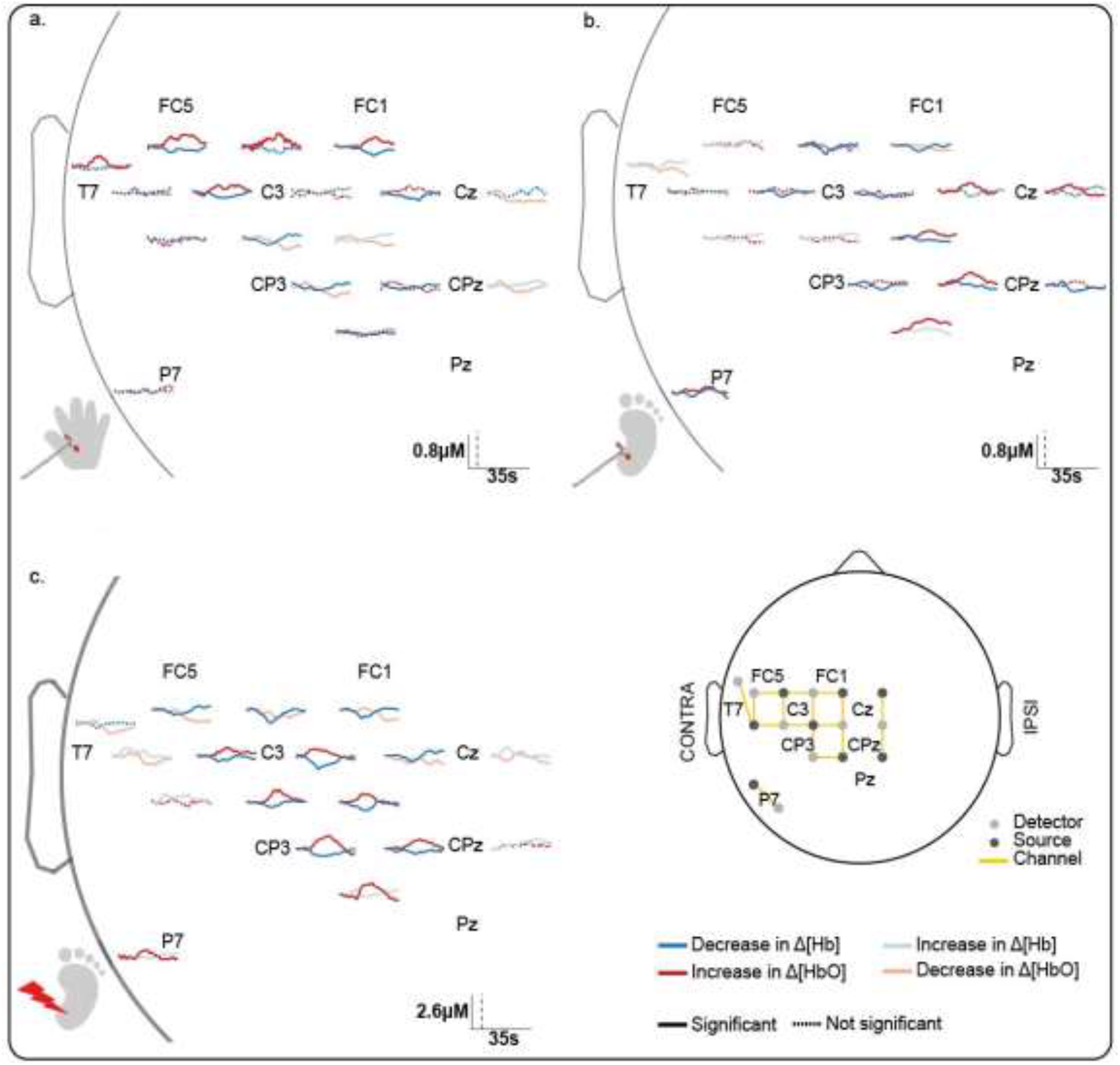
Channel-wise haemodynamic response following innocuous and noxious mechanical stimulation of hand and heel. Average Δ[HbO] (red) and Δ[Hb] (blue) duringN(a) hand touch (n=11), (b) heel touch (n=16) and (c) heel lance (n=11). Channels with significant increases in [HbO] and decreases in [Hb] (i.e. canonical response) during the activation period are shown with sold dark lines, inverse responses are shown with solid pale lines, and non-significant changes are shown with dotted lines. Black vertical line represents stimulus onset. Note the difference in the scale bar between touch and lance. For channels where a significant canonical and inverse response was found at different latencies, the canonical response only is depicted. (Details of individual channel responses are in **Figure 1 – Source Data 1**)

Image reconstruction of the channel data (Figure 2a and 2b) shows that the topography of touch activation in the newborn infant S1 is consistent with the known adult S1 topography: the area representing the foot lies in the superomedial postcentral gyrus, while the area for the hand is more inferior (Penfield and Boldrey, 1937; Harding-Forrester and Feldman, 2018; Willoughby et al., 2020).

**Figure 2.**
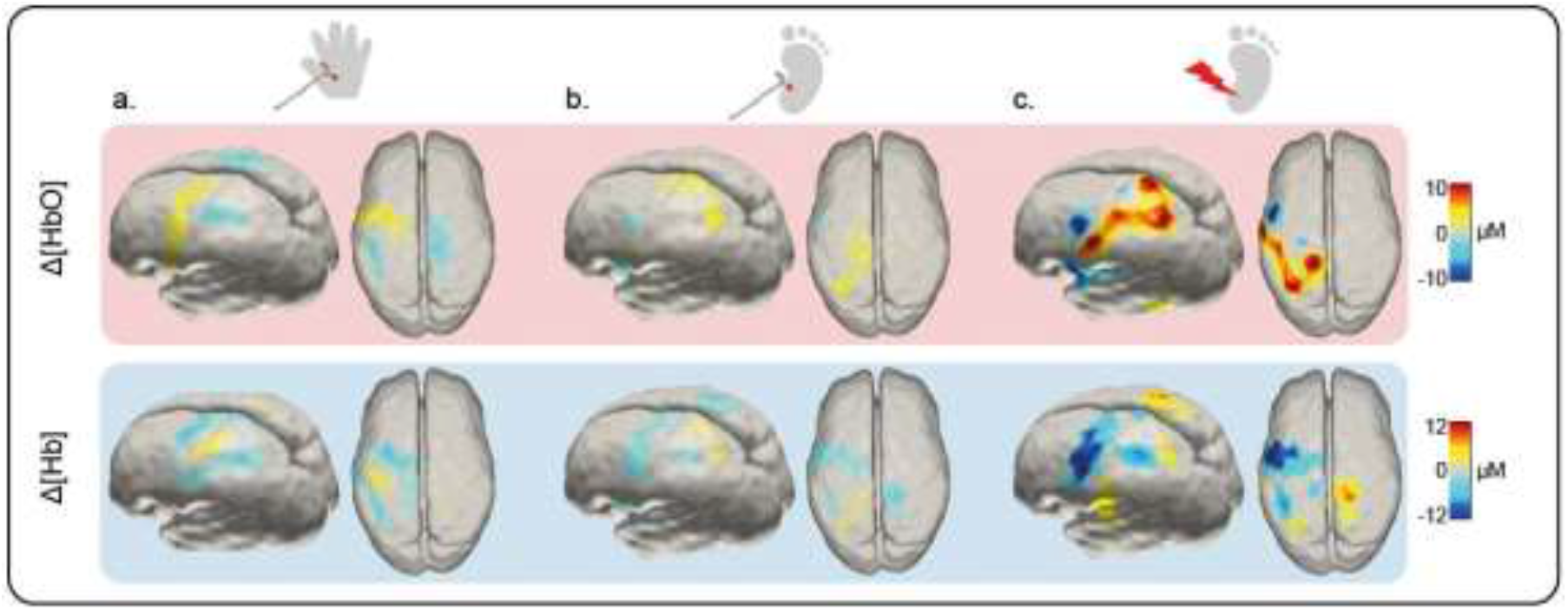
Image reconstruction at peak latency of the Δ[HbO] and Δ[Hb] response to an innocuous (touch) and noxious (lance) mechanical stimulation of hand and heel. Significant changes (compared to baseline) in Δ[HbO] (top row) and Δ[Hb] (bottom row) following (a) hand touch (n=11), (b) heel touch (n=16) and (c) heel lance (n=11).

### Noxious lance of the heel elicits widespread activation extending into inferior SI

We next mapped activation in the contralateral S1 following a noxious, clinically required, lance stimulus to the heel in newborn infants. The average channel response (**Figure 1c**) and the image reconstruction (**Figure 2c**) show the significant and widespread increase in [HbO] following lancing the heel, which extends beyond the somatotopic area for heel touch to encompass inferior areas of SI, which were associated with touch of the hand. Heel lance elicited significant increases in [HbO] in eight channels (including the control channel), with a maximum increase (0.96 μM at 14.5 s) at the channel corresponding to the CP2h 10:5 position. Four of the channels with a significant increase in [HbO] following the lance, also had a significant increase in [HbO] following touch of the foot. Notably one channel also displayed a significant increase following touch of the hand. The accompanying decrease in [Hb] was widespread (significant decreases in 11 channels; peak change 1.03 μM at 9.1 s), and an inverse response was found in all channels surrounding those with a canonical response (**Figure 1c and 2c**).

### Newborn infant nociceptive maps are not somatotopically aligned with touch maps

To test whether the widespread activity evoked by heel lance within S1 represents a true difference in the somatotopic mapping of touch and nociception, we compared the signal to noise, the individual channel activation and the overall spatial dimensions of haemodynamic activity evoked by the two stimulus modalities applied to the heel. Heel lance elicited a significantly larger increase in [HbO] and decrease in [Hb] compared to heel touch (maximum Δ[HbO]: 15.23 vs 4.13 μM, p=.007; maximum Δ[Hb]: -22.87 vs -4.34 μM, p<.001; **Figure 3b, Figure 3 – figure supplement 1**). However, the signal to noise of the lance heel activation was not higher than that of heel touch evoked activity (see Methods). Furthermore, the difference in amplitude between lance and touch was not accompanied by a difference in overall area of cortical activation evoked by the two stimuli (**Figure 3b and Figure 3 – Source Data 1**). Neither the location nor latency of peak activation differed between the two stimulus modalities (Δ [HbO]: distance between peaks = 1.65 mm, p=.776; difference in peak latency = 1.7 s [14.1 vs 15.8 s], p=.356; Δ [Hb]: distance between peaks = 0 mm, p=.957; difference in peak latency = 1.7 s [9.6 vs 11.3 s], p=.292). Only the spread of the peak Δ [HbO] was significantly larger following noxious heel lance (Δ [HbO] FWHM area: 66.63 vs 52.96 mm2, p=.015; Δ [Hb] FWHM area: 87.86 vs 83.10 mm2, p=.204). (Figure 3 – figure supplement 1). The key difference between the two patterns of activation is that the foot touch response is limited to the areas of the S1/M1 associated with the foot, whereas the lance response extends towards other more ventral regions of S1.

**Figure 3.**
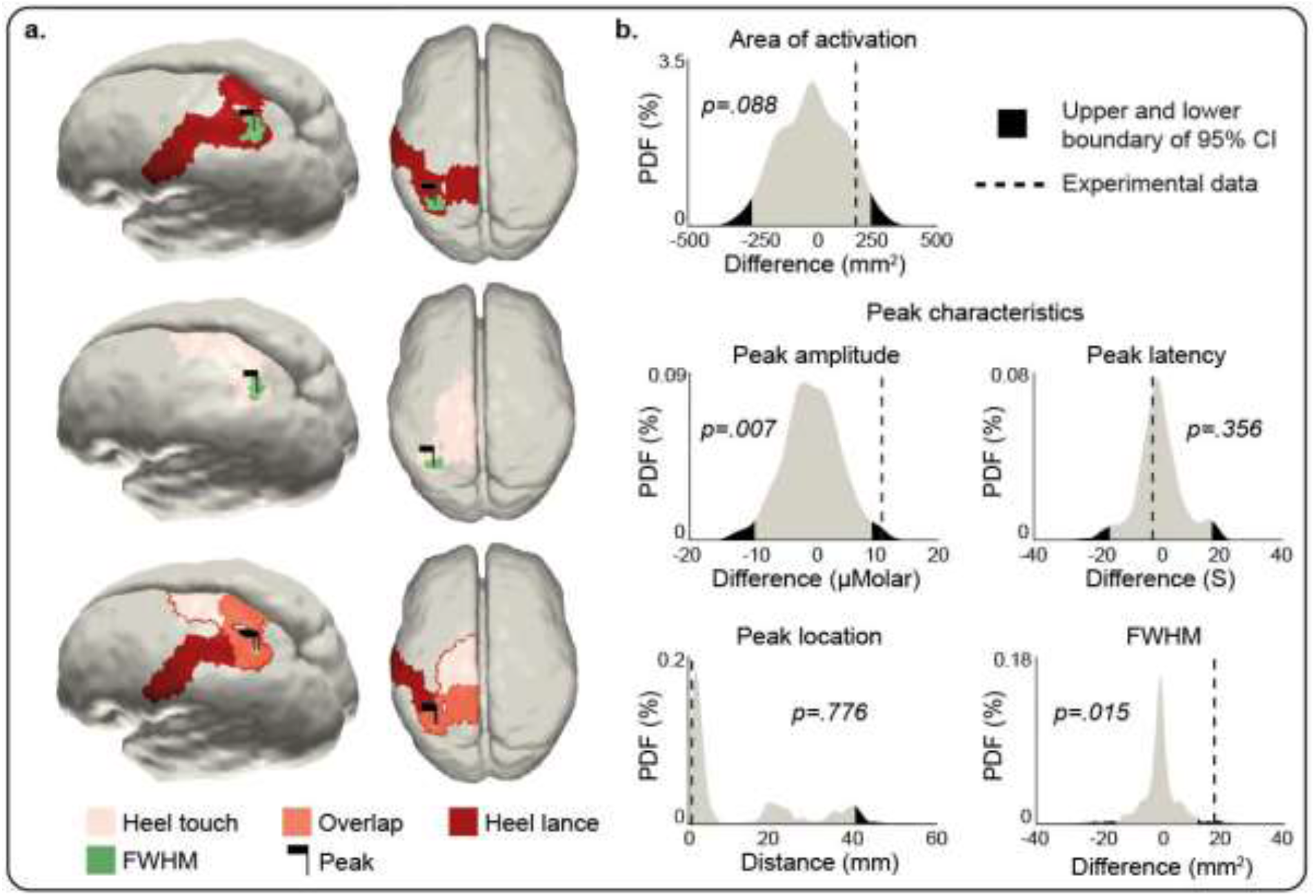
Comparison of the peak and area of activation of the Δ[HbO] response to an innocuous (touch) and noxious (lance) mechanical stimulation of the heel. (a) Area of significant Δ[HbO] changes following heel lance (red), heel touch (pink) and both (orange). Black flags demark the location of peak changes and green areas the extent of their full-width half-maximum (FWHM). (b) Statistical position of experimental differences between heel touch and lance in peak amplitude, FWHM, latency and location, and area of significant Δ[HbO] changes in respect to non-parametric null distributions obtained with bootstrapping and phase scrambling. The equivalent plots for Δ[Hb] are shown in **Figure 3 - figure supplement 1** and **Figure 3 - source data 1**.

**Figure 4.**
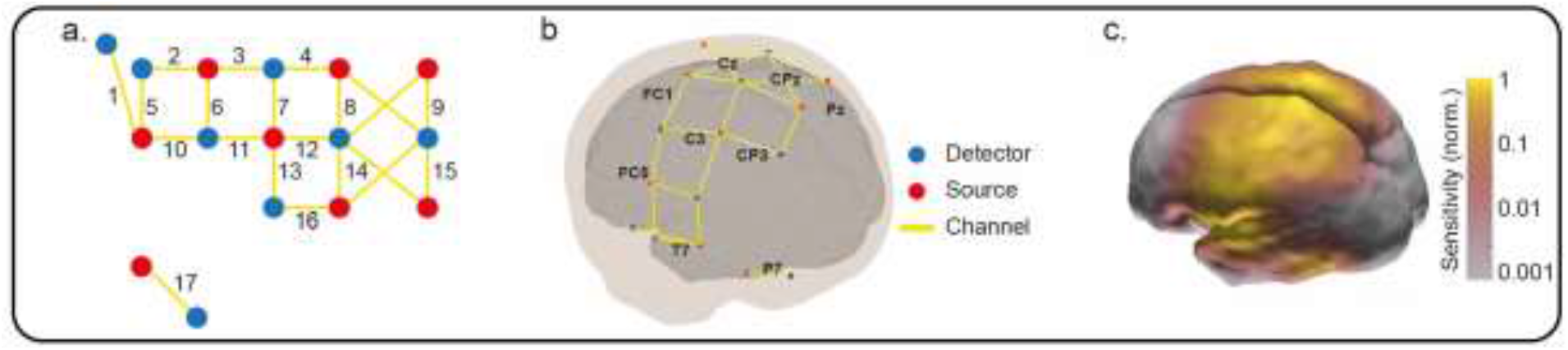
Optode locations and sensitivity map. (a) Channel reference numbers for Figure 1 – source data 1. (b) Locations of the fNIRS sources, detectors and resulting measurement channels registered to a 39-week anatomical atlas. (c) Normalized fNIRS sensitivity illustrating the spatial coverage provided by the channel arrangement in panel (b). This sensitivity map was calculated using the photon measurement density functions derived from the TOAST++ light transport modelling package.

## Discussion

### A widespread nociceptive topographic map in infant S1 that overlaps but is not aligned to the innocuous mechanoreceptive map

Somatosensory maps of cortical activity evoked by a cutaneous tactile or noxious stimulus provide a framework for localising the sense of touch or pain (Treede et al., 1999; Thivierge and Marcus, 2007). The adult primate S1 has a defined somatotopic organization of tactile and nociceptive cortical receptive fields (Andersson et al., 1997; Kenshalo et al., 2000) including spatially precise cortical maps of Aδ and Aβ afferent fibre input (Chen et al., 2011). Human fMRI studies show that adult somatotopic maps of noxious and non-noxious mechanical stimulation substantially overlap (Lui et al., 2008) and detailed analysis reveals a fine-grained somatotopy for nociceptive inputs in primary somatosensory cortex (S1) that are highly aligned with maps of innocuous tactile stimuli, suggesting comparable cortical representations for mechanoreceptive and nociceptive signals (Mancini et al., 2012). Here we have shown that this comparable representation is not present in the newborn infant S1 cortex.

Noxious mechanical stimulation evokes a larger peak increase in Δ[HbO] and decrease in Δ[Hb] compared to innocuous stimulation of the same body area at comparable location and latency as reported elsewhere (Bartocci et al., 2006; Slater et al., 2006; Verriotis et al., 2016b). The fact that the noxious activation is greater than the touch evoked activity (clearly seen because fNIRS provides scaled maps of the physiologically meaningful parameter, haemoglobin concentrations, unlike BOLD-fMRI) is presumably due to greater depolarisation and spike activity within the activated areas. However, it does not explain the differing topography reported here. The signal to noise ratio is not larger following noxious stimulation as we averaged data from repeated touches to compare with a single lance stimulus. Furthermore, the overall area of brain activity does not differ significantly between touch and lance of the heel, it is only that the areas of activation of the responses are not aligned. Within S1 itself, the infant has a distinct somatotopic map for touch, similar to that described in adults, with the area representing the foot lying in the superomedial postcentral gyrus, and the area for the hand located more inferiorly (Penfield and Boldrey, 1937; Blake et al., 2002; Akselrod et al., 2017), consistent with previous reports in newborn infants (Dall’Orso et al., 2018). Noxious heel lance, on the other hand, evokes a widespread activity within S1 , peaking in the same area of the superomedial postcentral gyrus as touch activity, but extending to the hand representation area. The multi-optode fNIRS array was placed over the contralateral perirolandic cortex and so the full extent of the nociceptive map is not known, but the data shows that the S1 somatotopic nociceptive map is not as precise as the touch map in the newborn.

### Measuring the cortical haemodynamic response to innocuous and noxious mechanical stimulation in neonates

fNIRS is ideally suited to a study of this kind as recording and sensory stimulation, including clinically required heel lance, can be performed at the infant cotside (Bartocci et al., 2006; Slater et al., 2010; Kashou et al., 2016; Verriotis et al., 2016b). Other methods of measuring this either do not provide sufficient spatial information and source localisation, such as EEG recording of nociceptive-related ERPs (Fabrizi et al., 2011; Jones et al., 2018) or are limited by the use experimental ‘pinprick’ stimulators, that for ethical reasons are not actually painful, such as in fMRI studies (Goksan et al., 2015).

The change in the Δ[HbO] and Δ[Hb] following sensory stimulation is a measure of neural activity: simultaneous vertex EEG and fNIRS recordings over S1 show that haemodynamic and neural responses are related in magnitude (Verriotis et al., 2016b). Following all stimuli, and consistent with the mature canonical response, channels showing a significant increase in Δ[HbO] also had a smaller decrease in Δ[Hb]. Regional overperfusion following neuronal activation, beyond that required by metabolic demands, means that less Hb is removed from the region compared to the oversupply of HbO. However, the decrease in Δ[Hb] was more widespread (but smaller in magnitude) compared to the localised increase in Δ[HbO]. This type of response, not previously reported in infants (de Roever et al., 2018), suggests that more blood is leaving the region (removing Hb) compared to the incoming supply (no significant change in [HbO] in peripheral channels) due to immature regulation of cerebral blood flow (CBF). There are multiple mechanisms by which blood vessels dilate and CBF increases following neural activation, including arterial CO_2_ and O_2_ concentrations, which relax/contract the smooth muscle cells of cerebral arteries and arterioles (Kety and Schmidt, 1948), and astrocyte and pericyte activity which contribute to vessel diameter and the propagation of vasodilation along the vascular tree (Takano et al., 2006; Cai et al., 2018), many of which are still developing in the newborn (Pryds and Greisen, 1989; Binmöller and Müller, 1992; Fujimoto, 1995) leading to rapid changes in CBF over the first postnatal days as cerebral circulation adapts (Meek et al., 1998).

These infants in this study were held skin to skin, swaddled in their mother’s arms in a naturalistic setting, which is a major advantage of nIRS recording over fMRI for human developmental studies of brain function. Video recording and investigator scoring confirmed that while some infant movement and maternal touching did take place and that some babies did move following the lance, these movements were varied in both body part and latency such that any associated cortical response would be removed during the averaging process. Furthermore, the chance of any larger movement from a few babies driving the widespread S1 response following the lance, is removed by the between group randomisation used to generate the null distribution of parameter differences. Finally, 33--50% of babies did grimace for up to 7 seconds following the lance, but if these facial movements mediated the response following the lance, this would have prolonged the peak or duration of the change in HbO, while in fact the latency and time course of the response to both stimuli was the same (see Figure 1).

### Differential development of somatosensory and nociceptive topographic maps

A whole-body topographical map of innocuous mechanical stimulation develops in the sensorimotor cortices over the early postnatal period in rats (Seelke et al., 2012), which represents the human final gestational trimester. In humans, distinct representations of the hands and feet can be observed from 31 weeks gestation, using fMRI (Dall’Orso et al., 2018), and from 28 weeks using neural activity recorded from the scalp (Donadio et al., 2018; Whitehead et al., 2018, 2019). The response to innocuous mechanical stimulation was more localised in S1 than the wider and less refined topographical map of noxious mechanical stimulation, suggesting a slower maturation of the S1 circuitry involved in nociceptive processing compared to touch processing in the infant brain.

In rodents, at every level of the developing somatosensory central nervous system, tactile processing matures before nociceptive processing (Fitzgerald, 2005; Koch and Fitzgerald, 2013; Chang et al., 2016, 2020; Verriotis et al., 2016a) consistent with a delayed refinement of a cortical nociceptive map. Widespread nociceptive cortical maps are consistent with infant pain behaviour, characterised by exaggerated and disorganised nociceptive reflexes in both rodent pups and human neonates (Fitzgerald, 2005, 2015), and which can fail to remove a body part from the source of pain (Waldenström et al., 2003). Nociceptive reflexes following noxious heel lance are larger in magnitude and significantly more prolonged in human infants compared to adults (Cornelissen et al., 2013) and have widespread cutaneous receptive fields that encompass the whole lower limb (Andrews and Fitzgerald, 1994). This lack of organisation could be reflected in the ascending spinothalamic and thalamocortical projections, delaying the maturation of S1 cortical nociceptive maps in the newborn. Topographic maps are established and aligned via multiple mechanisms, including molecular cues, spontaneous or sensory-dependent remodelling, and refinement. Initially, somatosensory maps are diffuse and overlapping, but in the rodent somatosensory cortex, excitatory thalamocortical afferents undergo activity-dependent refinement to sharpen these maps (Iwasato et al., 1997). Equally important is the maturation of inhibitory interneuron sensory maps which, in contrast, expand over development in an experience dependent manner (Quast et al., 2017). Slow developmental broadening of an inhibitory nociceptive network may explain the widespread nociceptive map in S1 and also the greater amplitude of EEG noxious responses in infants compared to adults (Fabrizi et al., 2016).

### Pain and the developing S1 cortex

This study highlights the importance of understanding the development of touch and pain processing in the human infant brain. The widespread S1 nociceptive topography discovered here implies that the infant S1 cortex would be unable to accurately localise noxious events and may lack the computational ability to reliably send noxious information to higher brain centres (Thivierge and Marcus, 2007; Harding-Forrester and Feldman, 2018). Heel lance is one of many skin-breaking procedures commonly performed in neonatal hospital care (Laudiano-Dray et al., 2020) and this study reveals the extent of cortical activation that follows just one such noxious procedure in the newborn. This contrasts with innocuous mechanical stimulation, such as touch, which activates a spatially restricted and somatotopically defined cortical area. Increasing evidence that repeated noxious experiences have adverse effects upon the developing brain (Ranger and Grunau, 2014; Duerden et al., 2018), underlines the importance of these results and the need for a better understanding of the mechanisms underlying the maturation of cortical nociceptive topographic maps.

## Materials and Methods

### Participants

Thirty-two infants (35-42 gestational weeks at birth, 0-7 days old, 12 female; **Table 1**) were recruited from the postnatal, special care, and high dependency wards within the neonatal unit at University College London Hospital. Infants received either 1) innocuous mechanical stimulation (touch) of the heel, 2) innocuous mechanical stimulation of the hand, or 3) a noxious mechanical stimulation (clinically required lance) of the heel. Six infants received touch stimulation of both the heel and hand. Similar high impact works using single trial noxious stimulation or multiple mechanical stimulations have yielded significant results with group sample sizes of 5-15 (Bartocci et al.,2006) and 10-15 (Arichi et al., 2012), respectively. Ethical approval for this study was given by the NHS Health Research Authority (London – Surrey Borders) and the study conformed to the standards set by the Declaration of Helsinki. Informed written parental consent was obtained before each study.

**Table 1.** Infant demographics. Demographic information about the subjects that received tactile and noxious stimuli of heel and hand.

|  | Heel lance | Heel touch | Hand touch | p |
| --- | --- | --- | --- | --- |
| **N** | 11 | 16 | 11 |  |
| **GA (weeks<sup>+days</sup>)** | 39<sup>+2</sup> (35<sup>+2</sup> – 41<sup>+5</sup>) | 39<sup>+4</sup> (35 – 42<sup>+3</sup>) | 39<sup>+2</sup> (37<sup>+5</sup> – 41<sup>+3</sup>) | .287 |
| **PNA (days)** | 4 (0 - 7) | 3 (0 - 6) | 3 (0 - 4) | .115 |
| **Females** | 4 (36%) | 6 (38%) | 5 (45%) | .889 |
| **Birth weight (g)** | 3134 (2220 - 4072) | 3250 (2360 - 4080) | 3300 (2450 - 3754) | .774 |
| **Caesarean deliveries** | 2 (18%) | 8 (50%) | 3 (27%) | .196 |
| **Head circumference (cm)** | 34 (32 – 35.5) | 34.25 (31 - 37) | 34 (32.5 - 36) | .900 |

*Values represent median and range or proportion. GA = gestational age (weeks from the first day of the mothers last menstrual cycle to birth); PNA = postnatal age (days since birth). No significant difference was found in any demographic parameter across the three groups (one-way ANOVA results in the last column).*

### Experimental design

Brain activity (fNIRS) was recorded following a clinically-required heel lance procedure or innocuous mechanical stimulation of the limbs at the bedside in the neonatal unit.

### Functional Near-Infrared Spectroscopy recording

Infants wore a 21-channel array consisting of 8 sources and 8 detectors with inter-optode distances of 2.5-4 cm. The array was secured over the pericentral area of the scalp on the side contralateral to the stimulation with a custom designed textile cap (EasyCap). The infants’ head circumference, ear to ear lateral semi circumference, and nasion to inion distance were measured and the cap was placed on the head by aligning specific 10/5 locations (Cz, T7). This optode arrangement provided sensitivity coverage for the whole somatomotor cortex contralateral to the stimulation site and of the medial part on the ipsilateral side (**Figure 4c**). One source-detector pair was placed at a more ventral posterior location of the scalp (P7 of the international 10/5 positioning system) (**Figure 4a**). This channel was sensitive to the posterior temporal lobe and worked as a control channel (**Figure 4b and 4c**). A continuous wave recording system was used with 2 wavelengths of source light at 780nm and 850nm and a sampling rate of 10Hz to measure changes in oxy-and deoxy-haemoglobin concentration (Gowerlabs NTS fNIRS system).

### Noxious mechanical stimulation

The noxious stimulus was a clinically required heel lance for blood sampling. Blade release was time-locked to the NIRS recording using an accelerometer attached to the lancet (Worley et al., 2012). The lancet was placed against the heel for at least 30s prior to the release of the blade. This was to obtain a baseline period free from other stimulation. The heel was then squeezed 30s after the release of the blade, again to ensure a post-stimulus period free from other stimuli. All lances were performed by the same trained nurse (MPL-D) using a disposable lancet, and standard hospital practice was followed at all times.

### Innocuous mechanical stimulation

Innocuous mechanical stimulation was delivered by light touch on the lateral edge of the infants’ palms and/or heels using a hand-held tendon hammer (ADInstruments). A piezo-electric sensor mounted on the hammer head provided a synchronising signal to the NIRS recording. A train of up to 15 touches (average = 11.5) was delivered to each limb with a variable inter-stimulus interval of 35 – 60 seconds. If the infant moved in the 30s pre-or post-stimulus the trial was removed (heel: average of 1.4 touches were removed in 8/16 infants; hand: average of 1.5 touches removed in 11/11 infants). This resulted in an average of 10.1 heel touches (range = 7 – 13) and 9.3 hand touches (range = 5 – 11) per infant.

### Recording infant movements

All infants were prone against their mother’s chest. The mother, who was inclined on a chair or bed, was instructed to avoid moving or stimulating the infant during the 1 minute before and after the release of the lance. Movements were minimized as infants were swaddled (wrapped securely in clothes/blankets) against the mother’s chest and the research nurse was holding the exposed foot throughout the period before and after the stimulus.

Infant movements, or changing tactile stimulation were recorded on video, which was synchronised with the NIRS recording using an LED light within the frame that was activated by the release of the lance (Worley et al., 2012). In case movements were obscured, a second researcher also recorded movements at the time of the study using a stopwatch. Movements or tactile stimulation were separated into body parts and coded per second as either 0 (not present) or 1 (present) for the 30 s post-stimulus. The total number of babies displaying each type of movement at each second post-lance can be seen in Table 2. Following the lance, 2 babies did not move, 6 babies made small movements (including: small or brief grimace, head nod, twitch, small hand movement), 4 babies made larger movements (including arms, large or prolonged grimace, nod of head), and 2 babies received tactile stimulation from the mother (including: positioning the head, stroking the face).

**Table 2.** Infant movements. The number of infants who displayed movements or received tactile stimulation from their mother each second in the 30s following lance.

| Post-lance<br>(s) | Number of infant and maternal movements |  |  |  |  |  | Mother touching face | Mother touching head |
| --- | --- | --- | --- | --- | --- | --- | --- | --- |
|  | Hand | Head | Face | Foot | Arm |  |  |  |
| 1 | 1 | 1 | 4 | 0 | 2 |  | 1 | 1 |
| 2 | 1 | 1 | 6 | 1 | 2 |  | 1 | 1 |
| 3 | 1 | 0 | 6 | 1 | 2 |  | 1 | 1 |
| 4 | 1 | 0 | 6 | 0 | 2 |  | 1 | 1 |
| 5 | 1 | 1 | 6 | 0 | 2 |  | 1 | 1 |
| 6 | 1 | 1 | 6 | 0 | 2 |  | 1 | 0 |
| 7 | 0 | 1 | 4 | 0 | 1 |  | 1 | 0 |
| 8 | 0 | 1 | 3 | 0 | 1 |  | 1 | 0 |
| 9 | 0 | 1 | 2 | 0 | 1 |  | 1 | 0 |
| 10 | 0 | 0 | 1 | 0 | 1 |  | 1 | 0 |
| 11 | 0 | 0 | 1 | 0 | 1 |  | 1 | 0 |
| 12 | 0 | 0 | 1 | 0 | 1 |  | 1 | 0 |
| 13 | 0 | 0 | 1 | 0 | 1 |  | 1 | 0 |
| 14 | 0 | 0 | 1 | 0 | 1 |  | 1 | 0 |
| 15 | 0 | 0 | 1 | 0 | 0 |  | 1 | 0 |
| 16 | 0 | 0 | 1 | 0 | 0 |  | 1 | 0 |
| 17 | 0 | 0 | 0 | 0 | 0 |  | 1 | 0 |
| 18 | 0 | 0 | 0 | 0 | 0 |  | 1 | 0 |
| 19 | 0 | 0 | 0 | 0 | 0 |  | 1 | 0 |
| 20 | 0 | 0 | 0 | 0 | 0 |  | 1 | 0 |
| 21 | 0 | 0 | 0 | 0 | 0 |  | 1 | 0 |
| 22 | 0 | 0 | 0 | 0 | 0 |  | 1 | 0 |
| 23 | 0 | 0 | 0 | 0 | 0 |  | 1 | 0 |
| 24 | 0 | 0 | 0 | 0 | 0 |  | 1 | 0 |
| 25 | 0 | 0 | 0 | 0 | 0 |  | 1 | 0 |
| 26 | 0 | 0 | 0 | 0 | 0 |  | 1 | 0 |
| 27 | 0 | 0 | 0 | 0 | 0 |  | 1 | 0 |
| 28 | 0 | 0 | 0 | 0 | 0 |  | 1 | 0 |
| 29 | 0 | 0 | 0 | 0 | 0 |  | 1 | 0 |
| 30 | 0 | 0 | 0 | 0 | 0 |  | 1 | 0 |

*Each movement or stimulation was scored as present (1) or not present (0) per infant, and the value in each cell represents the total number of infants (out of 11) for whom each movement or stimulation was observed.*

### Data pre-processing

All data were pre-processed in Homer2 (Huppert and Boas, 2005). Light intensity data were inspected for poor signal quality (signal intensity < 0.01, SNR < 2) resulting in 9 individual channels across lance trials (4%) being removed from further analysis, 12 across the heel touch trials (4%), and 6 across the hand touch trials (3%). Due to poor signal quality in the majority of trials, the 4 channels crossing over the midline were removed from all trials (**Figure 4a and 4b)**. Data were then converted into optical density, motion artefacts were detected (change in amplitude > 0.7 and/or change in standard deviation > 15 over a 1s time period) and then corrected using Wavelet filtering (Molavi and Dumont, 2012). Instrumental drift and cardiac artefact were removed with a 0.01-0.5 Hz bandpass filter. Optical density changes recorded from all channels (likely related to stimulus dependent systemic physiological changes) were removed using Principal Component Analysis ((Kozberg and Hillman, 2016; Tachtsidis and Scholkmann, 2016); 1 component removed). Finally, data were converted into changes in oxy-and deoxy-haemoglobin concentration (Δ[HbO] and Δ[Hb]) using the modified Beer–Lambert law (Delpy et al., 1988) with a differential path-length factor of 4.39 (Wyatt et al., 1990). The continuous signal was then epoched from -5 to 30 s around the noxious and somatosensory stimuli. Somatosensory stimuli were averaged for each subject.

### Signal to Noise

The signal to noise ratio (SNR) for lance and for touch were calculated. Despite the peak signal for lance being higher, the SNR is lower, because more touch trials were averaged in this study. The ratio of lance to touch SNR ((peak_lance_/peak_touch_)*(sqrt(n_lance_)/sqrt(n_touch_) = (0.96/0.30) * (sqrt(11)/sqrt(157)) = 0.85.

### Channel-wise data analysis

Pre-processed data were then averaged across subjects for each condition and analysed using custom MATLAB scripts (Mathworks; version 16b). For each channel, significant changes in Δ[HbO] and Δ[Hb] were identified with a two-tailed t-test (α = 0.01) comparing each time point post-stimulus against the baseline. This baseline distribution was calculated as the mean of the individual baselines (−5 – 0s before stimulus) according to:

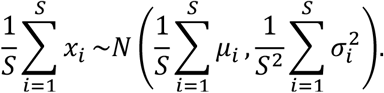

Where *S* is the number of subjects and 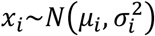 is the baseline for subject *i*. Bonferroni correction was used for multiple comparisons (17 channels x 300 samples = 5100 comparisons). Only changes (increases or decreases) which were continuously significant for at least 1 second (10% of the length of the post-lance period) were retained (Guthrie and Buchwald, 1991).

### Data analysis in image space

#### Image reconstruction

The channel-wise data was used to create functional images using a cortically constrained linear reconstruction approach. The fNIRS array was registered to a 39-week gestational age anatomical mesh model with 784391 nodes (Brigadoi et al., 2014) using tools from the AtlasViewer package (Aasted et al., 2015). Images were reconstructed using the DOT-HUB toolbox (www.github.com/DOT-HUB) and the TOAST++ light transport modelling package (Schweiger and Arridge, 2014) (www.github.com/toastpp), with zeroth-order Tikhonov regularization with a regularization hyperparameter of 0.1.

#### Assessment of changes in Δ[HbO] and Δ[Hb]

We first assessed the significance of the changes in Δ[HbO] and Δ[Hb] elicited by heel and hand touch and heel lance compared to baseline in image space. To do that, we reconstructed image time-series (i.e. one image reconstructed at each time-point) for each subject/stimulus. For each node, significant peak reconstructed changes in Δ[HbO] and Δ[Hb] were identified with a two-tailed t-test (α = 0.01) comparing the peak time point within a 5-second window around the peak latency derived from the channel-wise analysis against the baseline. As in channel-wise analysis this baseline distribution was calculated as the mean of the individual baselines (−5 – 0s pre-stimulus). Bonferroni correction was used for multiple comparisons (784391 nodes = 784391 comparisons). To display these results we: (1) reconstructed an image using the average channel-wise data within the 5-second window around the peak latency (averaged in time) for hand touch, heel touch and lance and (2) masked this image according to the result of the statistical test above.

#### Comparison of changes in Δ[HbO] and Δ[Hb] between heel lance and touch

Next, we wanted to compare the peak changes in Δ[HbO] and Δ[Hb] between the heel lance and touch conditions in image space. To do this, we calculated the difference in overall area of activation, and magnitude, latency, location, and spread of the peak change in Δ[HbO] and Δ[Hb], and compared these against a non-parametric null distribution.

Area of activation was defined as the cortical surface area with significant Δ[HbO] (or Δ[Hb]) changes. Difference in peak location was the Euclidean distance between the peaks. Peak spread was defined as the cortical surface area around the peak where changes in Δ[HbO] (or Δ[Hb]) were at least half of the peak change (full-width half-maximum, FWHM). Cortical surface areas (area of activation and peak spread) were defined by starting at the node with the peak change and continually expanding to include neighbouring nodes that were (1) connected by at least 2/3 face edges, and (2) had a significant change from baseline.

The non-parametric null distribution was derived by calculating these differences between randomly selected sets of surrogate image time-series (bootstrapping on surrogate data). We here describe how we obtained surrogate image time-series and then how we conducted bootstrapping.

Each individual recording (i.e image time-series) can be considered as the linear sum of a signal of interest (i.e. the response to the stimulus) and a stationary random noise component. The assumption is that the signal is the same in each recording while the noise changes. Therefore, if we were to conduct another recording on another subject the new data would be the linear sum of the *same* signal that we find in the original data but *different* random noise. Creating surrogate data consists in generating new random noise to add to the signal estimated from our data. To do this we: (1) estimate the signal by averaging across individual recordings (i.e. subjects) in response to the same stimulus modality; (2) isolate the noise in our data by subtracting this estimate from each recording; (3) *phase-randomise* each noise time-series. Phase-randomization is applied independently to each node time-series in the frequency-domain. This means that the phase component of the complex-valued signal is rotated at each frequency by an independent random variable chosen from the uniformly distributed range of 0 and 2π (Theiler et al., 1992). At the end of this process we have a new set of surrogate noise time-series.

To generate the full non-parametric null distribution against which to compare our data, we used bootstrapping. To estimate each sample of the null distribution, we calculated the differences in area of activation and peak amplitude, latency, position and FWHM between two random sets of surrogate data without any systematic difference. To create the random sets, we: (1) pooled together all the newly obtained surrogate noise time-series; (2) added the grand average (across lance and touch) signal (as we do not want systematic differences between sets to estimate a null distribution); (3) randomly split (with repetition) these surrogate data into two sets. We repeated this 1000 times in order to obtain the full non-parametric null distribution (bootstrapping). An experimental difference outside the 95% confidence interval was considered significant (p < 0.05).

### Data sharing

All raw data files are open access and are available to download from Figshare (https://doi.org/10.6084/m9.figshare.13252388.v2).

## Acknowledgments

This work was funded by the Medical Research Council UK (MR/M006468/1, MR/L019248/1, and MR/S003207/1). RJC is funded by EPSRC Fellowship EP/N025946/1

## Supplements and Source Data

**Figure 1 – Source Data 1.**
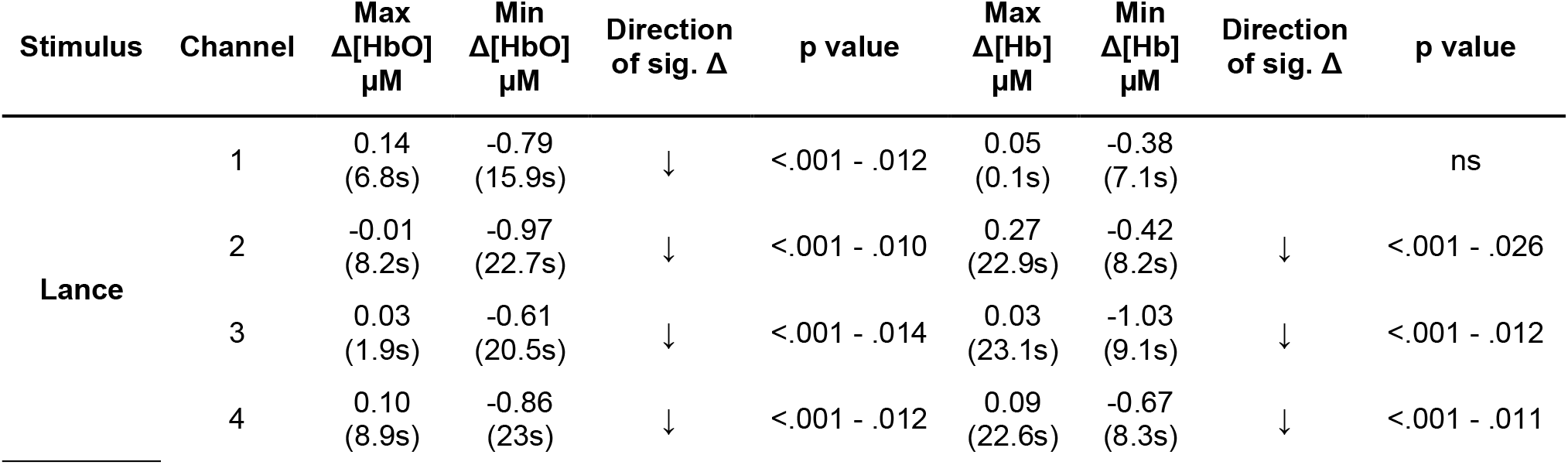
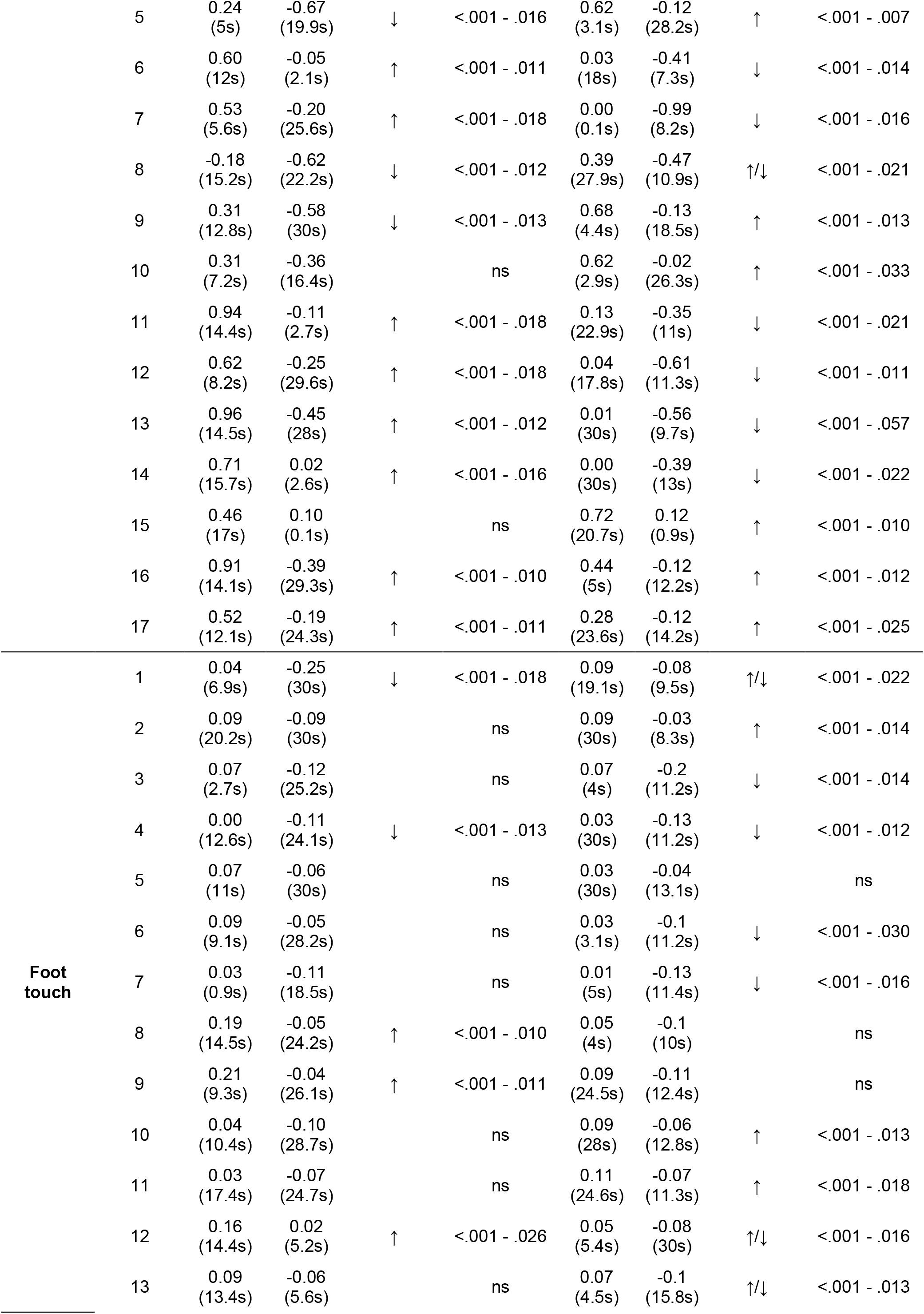
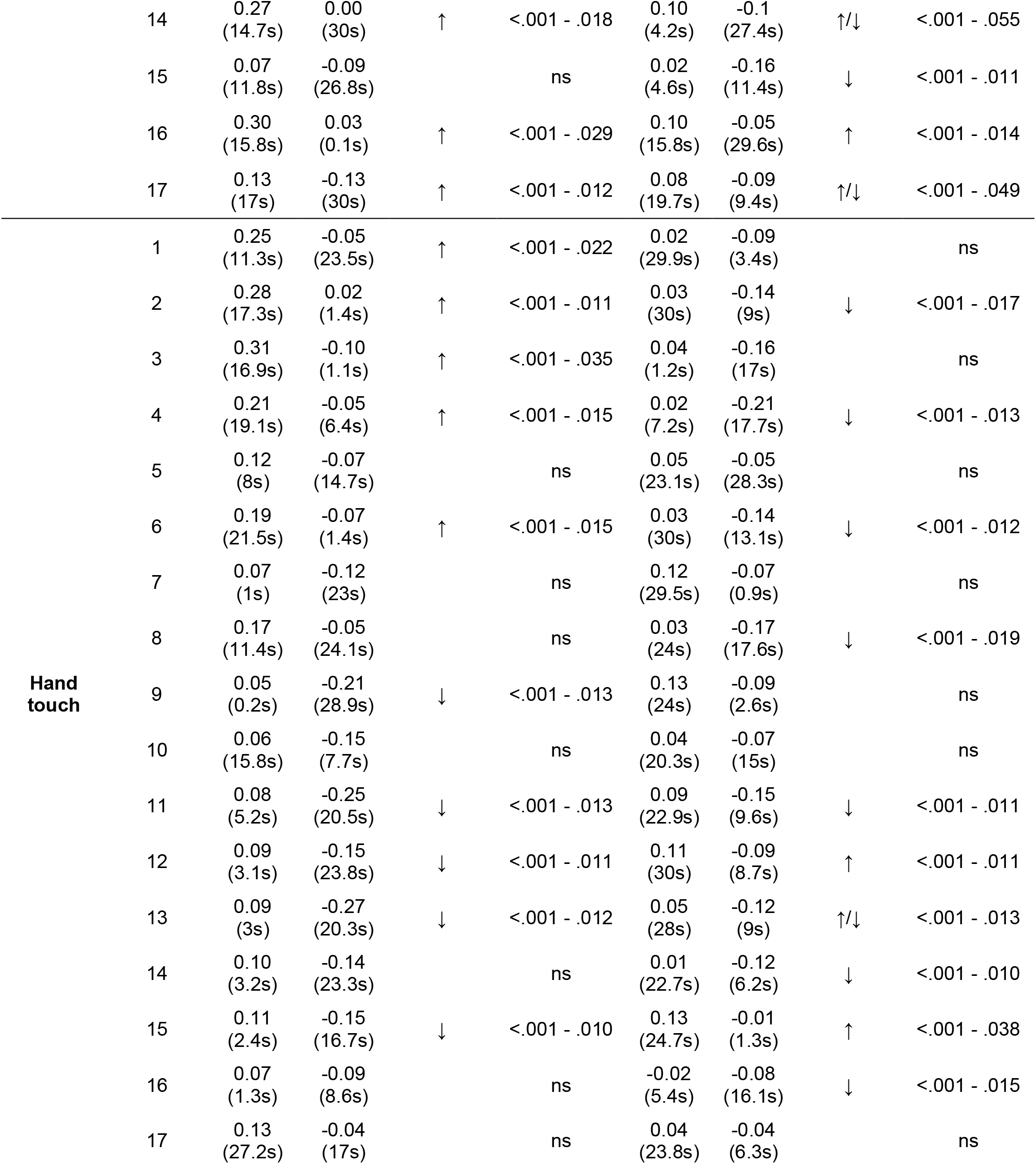
Significant concentration changes at each channel following innocuous mechanical stimulation (touch) of the heel and the hand and following heel lance. Minimum and maximum Δ[HbO] and Δ[Hb] at every channel and the corresponding latency. The range of Bonferroni-corrected p values across all significant timepoints is provided. ↓ = significant decrease only, ↑ significant increase only, ↑/↓ = both a significant increase and decrease were observed at different latencies. The location of each channel is shown in Figure 4.

**Figure 3 – figure supplement 1.**
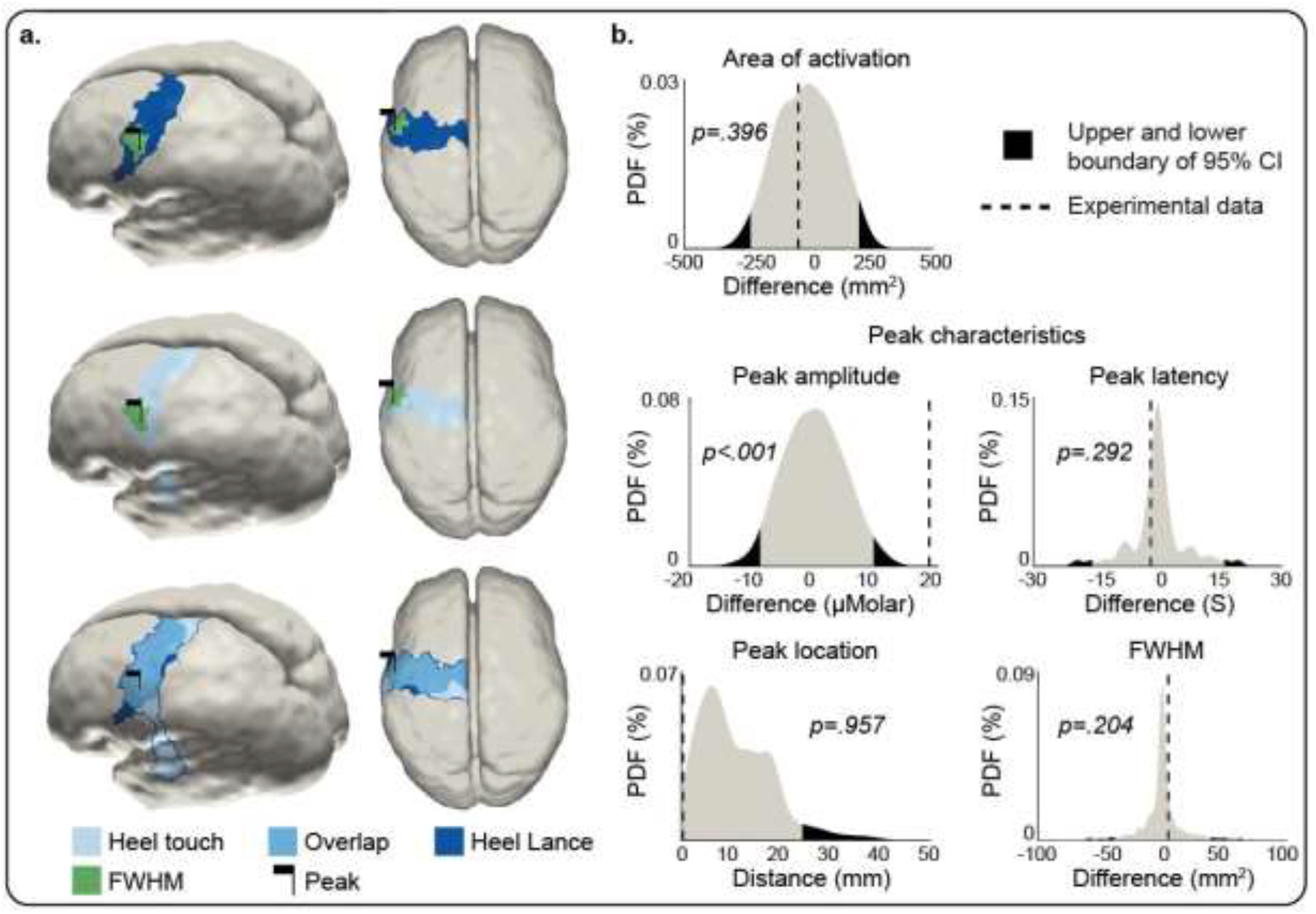
Average Δ[Hb] during a heel lance and innocuous mechanical stimulation (touch) of the heel and comparison of the peak change. Left (a): Average Δ[Hb] following a heel lance and innocuous mechanical heel stimulation at the peak latency; Dark blue (lance) and pale blue (touch) patches represent the cluster of neighbouring nodes which are significantly larger (HbO) than baseline and mid blue is the area of overlap. Green patches represent the full width half maximum (FWHM) within these clusters. Black flags denote the location of the peak change. Right (b): Null distribution of the differences between amplitude, latency, location, and spread of the peak change, obtained with bootstrapping and phase scrambling (1000 iterations). Black shaded areas represent the 2.5 and 97.5 (or 95 for location) percentile of the distribution and black dashed lines represent the values obtained with the experimental data. The equivalent plots for Δ[Hb] are shown in Figure 3 – figure supplement 1 and Figure 3 – source data 1.

**Figure 3 – Source Data 1.**
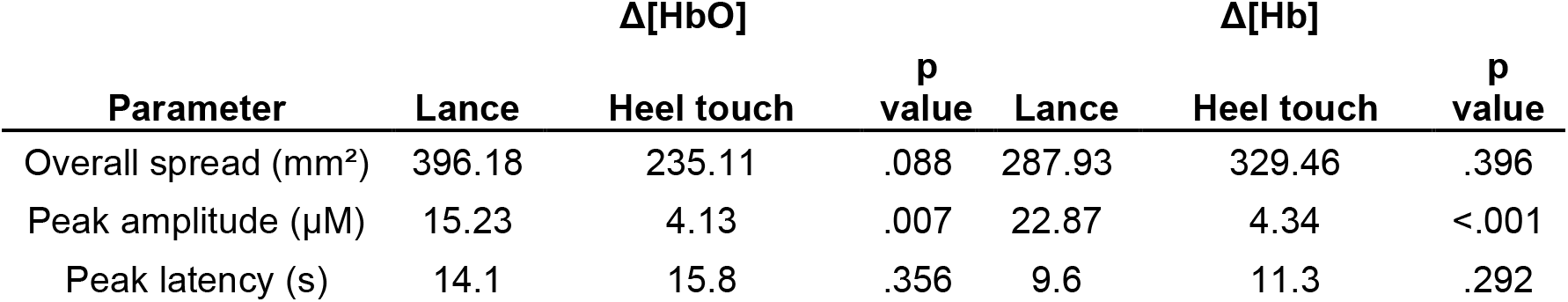
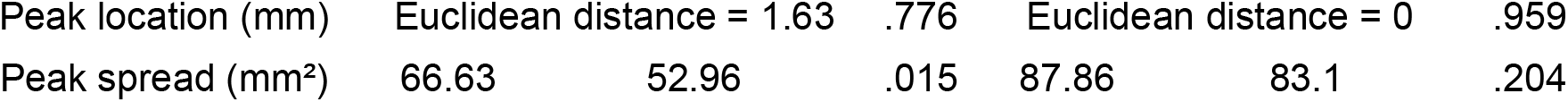
Comparison of Δ[HbO] and Δ[Hb] between heel lance and heel touch Parameter estimates for the Δ[HbO] and Δ[Hb] following a heel lance and heel touch, and p value from statistical comparison. The Euclidean distance between the heel touch and heel lance peak locations has been provided rather than x,y,z coordinates of each peak. Linked to Figure 3 and Supplementary Figure 1.

